# Expanding the Synthome: Generating and integrating novel reactions into retrosynthesis tools

**DOI:** 10.1101/2025.03.07.642029

**Authors:** Maranga Mokaya, Charlotte M. Deane, Anthony R. Bradley

## Abstract

Synthesising compounds is a major component of the time and cost of drug discovery. Retrosynthesis approaches have grown in prominence in efficiently predicting synthetic routes. For such tools to be truly impactful they should be able to effectively incorporate rarely seen chemical reactions and synthesise unknown molecules. We show that whilst existing tools can often effectively replicate known routes in familiar chemical spaces, they tend to fail in less explored regions. To overcome this challenge, we present a novel approach that integrates under-utilised reactions, resulting in improved performance in nearly 70% of tested cases. We introduce a tool that identifies, prioritises, and incorporates entirely new chemical transformations, demonstrating an average reduction in predicted synthesis costs of ∼ 20%. Importantly, our tool can highlight new chemical reactions that should be prioritised for development to improve synthesis efficiency. Our findings lay the groundwork for next generation retrosynthesis tools, which will be capable of driving the discovery of novel chemical transformations and more efficient exploration of chemical space.

## Introduction

Drug discovery can be considered a search problem. To generate therapeutic agents with desired properties it is necessary to traverse chemical space (i.e., all possible chemical structures) to synthesise, and test molecules. The challenge is not merely in identifying molecules with the desired biological properties but also finding those that are safe and affordable. The drug discovery process is expensive and inefficient. Each new drug is estimated to cost between $0.94 and $2.8 billion with a median development time of 8.3 years.^1–3^ Therefore, any tool that improves this process will have significant impacts on the diseases we can target and the populations that can benefit.

Computer-aided solutions to improve efficiency have increased in prominence within the drug discovery process in areas such as molecular generation, property prediction, and synthesis.^4–12^ Retrosynthesis tools have been created to help the elucidation of efficient pathways to a target molecule.^13^ Retrosynthesis is the process of deconstructing a target molecule into its precursors by applying the reverse of a forward chemical reaction repeatedly. Early retrosynthesis tools relied on hard coding of synthesis rules and heuristics.^14,15^ Current methods using deep learning and reaction databases^16–18^ have enabled more comprehensive searches for appropriate synthetic routes to target molecules.^13^

Multi-step retrosynthesis tools typically comprise three essential components. First, a single-step reaction prediction model, second, a search algorithm and third, a building block library. The single-step reaction prediction model determines both the type and location of potential reactions, generating sets of viable reactants for a given target molecule or intermediate.^16–21^ These models employ various approaches to molecular representation, synthesis rule extraction and reagent generation. To enable multi-step pathways, search algorithms compare and rank multiple reaction sequences to identify optimal synthetic routes. Whilst any tree search algorithm could theoretically be used, Monte-Carlo tree search^22,23^ and heuristic A* search^24^ are the most popular methods. The building block library typically consists of a curated set of purchasable molecules from which the target molecule can be synthesised.

Reaction prediction models can be categorised as either template-based or template-free determined by their single-step model approach. Template-based systems require an additional component: a template library comprising of reaction templates from which the single-step model selects the most appropriate reaction. For instance, Segler *et al*. developed a neural network that prioritises reaction rules based on a target molecule’s fingerprint representation.^25^ Template-free approaches operate by generating reagents without predefined reaction templates. Karpov *et al*. demonstrated this approach with a text-based model that translates target SMILES directly into reagent SMILES using a transformer model.^26^ Both methodologies have been successfully integrated into multi-step retrosynthesis tools to generate complete synthetic routes.^27–29^

In our study, we focus on two types of retrosynthesis tools: First, we utilise the template-based variant of the popular AiZynthFinder framework which incorporates a central neural network to guide the search process. At each iteration, this network recommends the most suitable template from a predefined library to further breakdown the target molecule. We also use Postera’s Manifold^30^ which adopts a template-free approach. Based on the molecular transformer^31^, it regressively generates reagents given an input target SMILES. This process is repeated till search conditions are exceeded or a set of purchasable precursors are identified.

Recent developments in synthesis prediction models have mostly focused on improving the accuracy or validity of suggested routes. For example, incorporating better molecular representations or experience-based model training. Alongside improved datasets and more standard benchmarks.^19,36–41^

Synthesis prediction models have been shown to efficiently generate synthetic pathways, they also face some limitations. Notably, the performance metrics used to benchmark these tools reward the replication of existing reaction data, biasing results toward established synthetic preferences. This approach may be suboptimal, as studies have demonstrated that chemists typically employ a limited set of transformations, thereby sampling only a fraction of potential reaction space and limiting accessible chemical space.^42^

We show that current tools struggle to solve synthetic routes that require under-utilised transformations. We propose that expanding accessible chemical space demands more innovative tools that can capture and incorporate more creative chemistry where necessary.

To overcome this limitation, we propose a new tool, RetroMix (see Code availability) which builds on the AiZynthFinder framework. This tool provides a novel method to incorporate multiple synthesis strategies, including novel chemistry, into a template-based system. We show how, in addition to solving target molecules with higher success and lower cost, our method can highlight and prioritise under-utilised and novel transformations that should be developed. This hybrid approach enhances retrosynthesis performance, enabling exploration of broader chemical spaces and suggests methods for improving the synthesis of complex molecules.

## Results & Discussion

Modern retrosynthetic tools are powerful techniques for finding synthesis pathways.^41^ While these tools have been shown to often supply a quick and simplified approach to solve synthesis pathways, in this work we investigate how effectively current tools explore reaction space.

### Retrosynthesis Performance on Popular Targets

We investigated how well retrosynthetic tools navigate reaction space to solve molecules from well-explored and less-explored areas of chemical space. To do this we evaluated the performance of a template-based (AiZynthFinder) and a template-free (Postera) retrosynthesis tool on sets of actives against two classes of targets.

A popular method for benchmarking retrosynthesis tools is how well they can generate known routes for a set of target molecules.^40,43^ This practice, while valuable in established chemical spaces, poses challenges when applied to under-explored areas. We know that the reactions and building blocks available are heavily biased to those that have proven most effective thus far^25^. We suspect these biases will reduce the tools performance on less well explored molecules. To assess this, we attempted to find routes for molecules known to be active against kinase and non-kinase targets. Kinases are a popular class of drug target. Therefore, their actives present a well-studied and explored area of chemical space.^44^ Conversely, less prevalent non-kinase targets are likely to be less well studied, and their actives are likely to be in more remote areas of chemical space.

Table 1 reveals a significant disparity in the performance of retrosynthesis tools between molecules active against kinase and non-kinase targets. AiZynthFinder and Postera exhibited higher success rates for kinase targets (67% and 94%, respectively), demonstrating their proficiency in well-documented chemical spaces. However, both tools showed reduced performance for non-kinase targets (58% and 82% success rates), suggesting limitations in less explored chemical spaces. Analysis of the overlap between solved and unsolved molecules across target classes reinforced this trend. For kinase actives, 65% were solved by both tools, compared to only 55% for non-kinase actives. Moreover, a substantially higher proportion of non-kinase targets (17%) remained unsolved by both tools, compared to only 5% of kinase actives. Further analysis of molecular properties revealed substantial overlap between kinase and non-kinase targets (see S1), suggesting that the observed disparity in model performance is not due to a divergence in molecular properties. These findings support our expectation that current retrosynthesis methods perform more effectively on well-studied molecules but struggle in less well explored areas of chemical space.

**Table 1:**
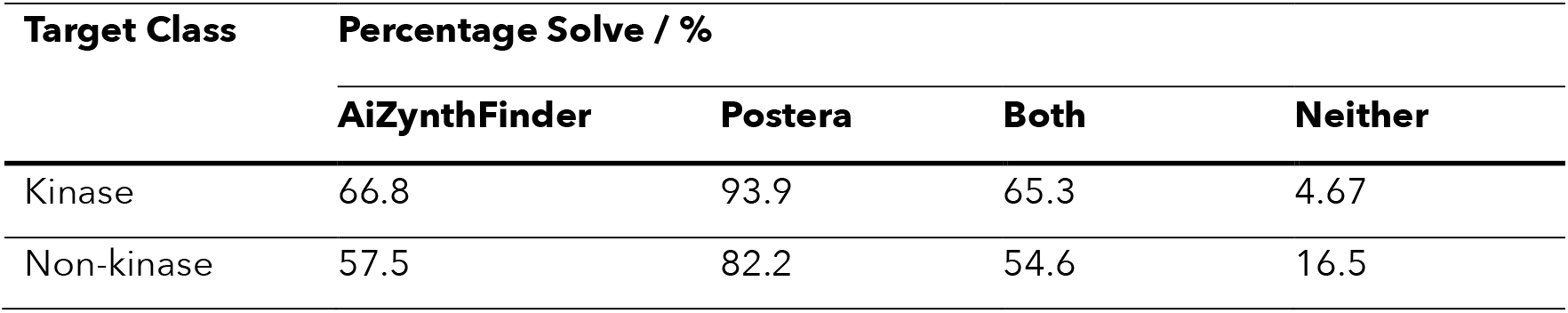
Performance of AiZynthFinder and Postera on a set of kinase and non-kinase actives. Performance for both tools was better on kinase targets relative to non-kinase targets. Both and neither shows the overlap in solved and unsolved molecules for both methods.

| Target Class | Percentage Solve / % |  |  |  |
| --- | --- | --- | --- | --- |
|  | AiZynthFinder | Postera | Both | Neither |
| Kinase | 66.8 | 93.9 | 65.3 | 4.67 |
| Non-kinase | 57.5 | 82.2 | 54.6 | 16.5 |

Comparing the two methods, the superior performance of Postera is unsurprising, given its freedom from template library restrictions. Its translation-based training regime offers greater flexibility, enabling more creative solutions where template-based methods may struggle. Particularly in resolving unusual intermediates generated by less prevalent templates. Further to this, we observed similar results when comparing the template-free and template-based variants of AiZynthFinder (see S2). Postera’s performance may partly stem from incorporating infeasible reactions. These should be viewed as potential avenues for improving synthesis rather than guaranteed routes. Despite its ability to include novel and potentially infeasible reactions, Postera’s performance still falters when route searching for less well-documented molecules.

Understanding the possible causes of failure is important for understanding a retrosynthesis tool’s limitations. We explored the causes of failure and found that template and building block availability place limits on retrosynthesis performance (see S3 for more details). We posit that the reduced performance for non-kinase actives is related to with their popularity in research and therapeutic development. The lack of patented reactions and tailored building block libraries for less common targets contributes to the ineffectiveness of retrosynthetic tools for less explored molecules.

### Library Frequency and Template Use

Our findings highlight how the challenges faced by retrosynthetic tools in less studied areas of chemical space are potentially rooted in biases within reaction libraries. To examine the strength of this hypothesis, we examined the relationship between template usage in solved routes and the occurrence of these templates in the US Patent Office (USPTO) library of reactions. ^45,46^

Figure 1 shows that on average the more a template is used, the more often that transformation appears in the USPTO template library. The Pearson correlation coefficient between template use and library occurrence was found to be *r*(3746) = 0.599 (p-value < 0.001) confirming a significant correlation. We also see more templates with low USPTO occurrences used more often for unsolved routes (see S4). This follows from our earlier results; the more complex, and less well-documented molecules require less popular templates. Once these templates are employed during a search, we postulate that the tools struggle to resolve the subsequent intermediates, leading to failed route generation attempts.

**Figure 1.**
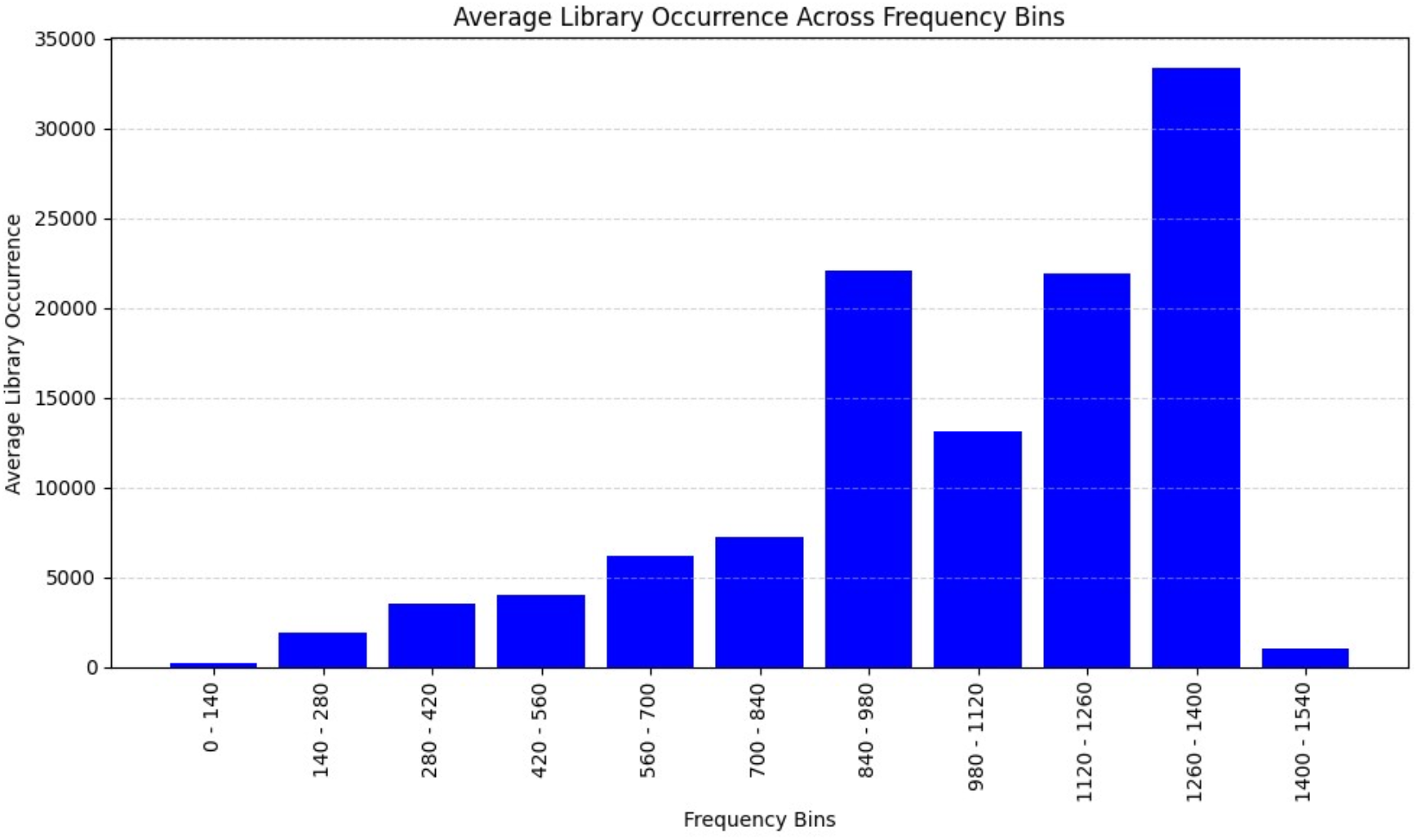
Relationship between template usage and library occurrence in the USPTO database. The more often a template is used, the more often it appears in template libraries. Each template was grouped according to its frequency across all molecules. Then the average library occurrence for each bin was determined.

Finally, we investigated how many templates were used by AiZynthFinder and Postera across the molecules that were attempted by both. AiZynthFinder successfully used 49.3% of the templates used by Postera, whereas Postera used only 29.0% of templates used by AiZynthFinder. This highlights the distinct strategies used by each method and suggests that the exploration exhibited by Postera is a contributor to its superior route-finding performance.

### Analysis of Reaction Space Exploration

The results above suggest that template-based and template-free methods can struggle (to varying degrees) to resolve routes for less well-documented molecules due to the need to use less prevalent transformations. To investigate this further, we analysed the diversity and distribution of templates in solved and unsolved kinase active routes. We quantified the number of distinct template clusters at each reaction step by applying the DBSCAN clustering algorithm^47^ to reaction fingerprints of each template used. A tool unable to explore reaction space well may try to use templates outside of the popular set leading to incomplete synthesis pathways for unsolved molecules. Conversely, we expect that a tool able to resolve a diverse set of routes would exhibit a comparable spread of templates for both solved and unsolved molecules. In this case any failures are more likely to be due to gaps in available building blocks.

Figure 2 shows the number of distinct template clusters at each step of solved and unsolved synthesis pathways. These results show that, firstly, unsolved routes tend to have more steps (16 for unsolved compared to 14 for solved), indicating more extensive branching to solve a difficult route within the search constraints. Secondly, for unsolved routes, the number of distinct template clusters remains higher for longer compared to solved routes (from step 3 onwards, there are more template clusters for unsolved routes). This indicates a larger diversity of templates are used in the routes that ultimately fail. These failures may stem from the increased diversity of generated intermediate molecules, which require a more varied set of templates including less popular ones for resolution. These tools are not optimised to use these templates and therefore struggle to do so within the search constraints.

**Figure 2.**
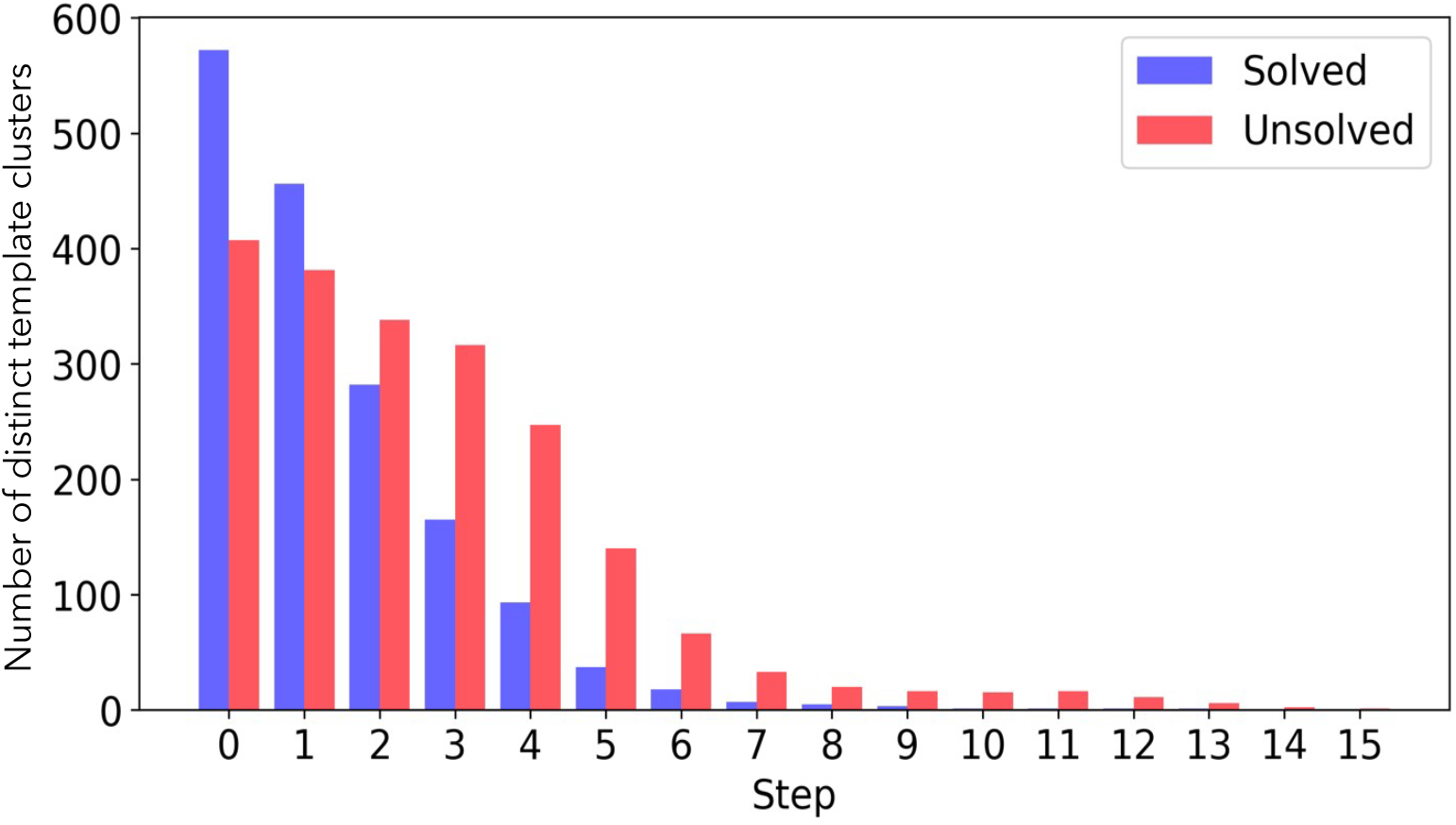
Diversity of solved routes falls quicker through a pathway compared to unsolved routes. This figure illustrates the number of distinct template clusters at each stage of a synthesis pathway for both solved and unsolved kinase actives using AiZynthFinder. Initially, solved routes exhibit greater diversity, but by the third step, unsolved routes explore a broader set of templates, resulting in unsuccessful outcomes

### Comparing popular or overlooked strategies to improve retrosynthesis performance

So far we have shown that retrosynthesis tools struggle to solve molecules in underexplored chemical space. We investigated ways to leverage the differences in search strategies between template-based and template-free methods to better explore chemical space and identify more efficient synthesis pathways. Additionally, we sought to understand the impact of introducing underutilised transformations into route searches, and from this determine which transformations should be optimised and more widely incorporated into synthesis strategies.

We collected data on 80 kinase and 80 non-kinase targets (see S5) then solved routes for 20 active molecules per target using AiZynthFinder and Postera. We identified the most frequently used reactions for each method to define their respective strategies. Template categories are detailed in Table 2 in addition to Novel templates which will be explained in the next section.

**Table 2:** Optimised template classification. Descriptions of the three template classes used in each optimisation strategy. Popular and overlooked templates are in the template-based (TB) library, whereas novel templates are not. These templates are generated by template-free (TF) tools.

| Class | Definition | Significance |
| --- | --- | --- |
| Popular | Frequently used in cheapest TB routes | Well-documented, established reactions |
| Overlooked | In TB library, unused in TB search, used in TF search | Under-utilised documented reactions |
| Novel | Not used by TB, not in TB libraries, applied by TF | Undocumented chemistry |

Popular templates from AiZynthFinder are well documented in synthesis data (see Figure 1). These templates are used in the cheapest routes for each solved molecule and are ranked by frequency (see methods, template classification). Overlooked templates characterise successful synthesis strategies accessible to AiZynthFinder but not used. These are Postera’s most frequently used templates in its cheapest routes which were absent from AiZynthFinders’ routes.

We conducted two experiments: first, a standard route search, followed by a repeated search for each active molecule with template optimisation for each strategy, using route price as the guiding metric. Our optimisation process (see methods) simulates the introduction and promotion of specific templates in actual synthesis, making them more accessible and practical for real-world applications.

We developed the Post Optimisation Score (POS, Eqn. 1), to quantify the effects of optimising specific reactions. This metric balances the cost change for synthesising each molecule (*D*), the cost associated with newly solved molecules (*E*), and the total number of solved molecules (S), with each term being adjustable through their coefficient (*α, β* and *γ*) to reflect desired outcomes. *α* is defines the weight of the cost change score after optimisation, *β* defines the weight of newly solved molecule cost score and *γ* defines the weight of the solved molecule score. For instance, if maximising the total number of solved molecules is the primary objective, the weight of *γ* should be increased. Conversely, if reducing synthesis costs is the goal, the weight of *α* should be increased. Our scoring method shows the practical implications that transformation optimisation may have on objectives important to chemists.

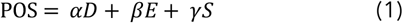

Post-optimisation score (POS) is a metric to describe the effect of optimising specific reaction on retrosynthesis performance (1). *D* is the cost change for synthesising each molecule after optimisation (2). E is the cost associated with newly solved molecules relative to the mean of previously solved molecules (3). *S* is the total number of solved molecules (4). *α, β* and *γ* are coefficients that can be altered to highlight the three different performance metrics. In this experiment, the route price is a function of the price of all purchasable precursors, the sum of precursors and reaction steps (see methods, performance evaluation).

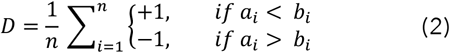

*D* is the difference in costs of molecules solved before and after optimisation. *n* is the number of molecules solved by both. *a*_*i*_ is the cost of synthesising a molecule after optimisation, *b*_*i*_ is the cost before optimisation.

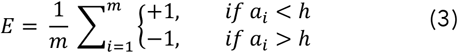

*E* is the difference in cost between newly solved molecules after optimisation and the average cost of previously solved molecules. *h* is the mean cost of previously solved molecules and m is the number of newly solved molecules.

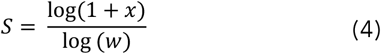

S scores how many more molecules are solved after optimisation.

*x i*s the number of extra molecules solved after optimisation and w is the total number of molecules for which synthesis pathways were attempted for this target.

*Equation 1: Mathematical description of Post Optimisation Score (POS) and its components. POS is a quantitive description of that retrosynthesis performance after transformation optimisation*.

In Figure 3a, we present the solvability-prioritised POS scores (0.3, 0.2 and 0.5 for *α, β* and *γ* respectively) for 80 kinase targets. This choice of parameters gives a focus on the overall number of molecules solved after optimisation and is an example of a common goal for medicinal chemists (see S6 for alternative POS results with different goals). We found that promoting Popular templates often has a more significant positive effect than introducing new ones for kinase targets. Nonetheless, overlooked strategies can also yield substantial benefits as we see several examples where Overlooked template optimisation has a greater positive effect than Popular template optimisation. We observe positive POS scores for Popular template optimisation in 89% of targets and for Overlooked strategies in 69% of targets.

**Figure 3.**
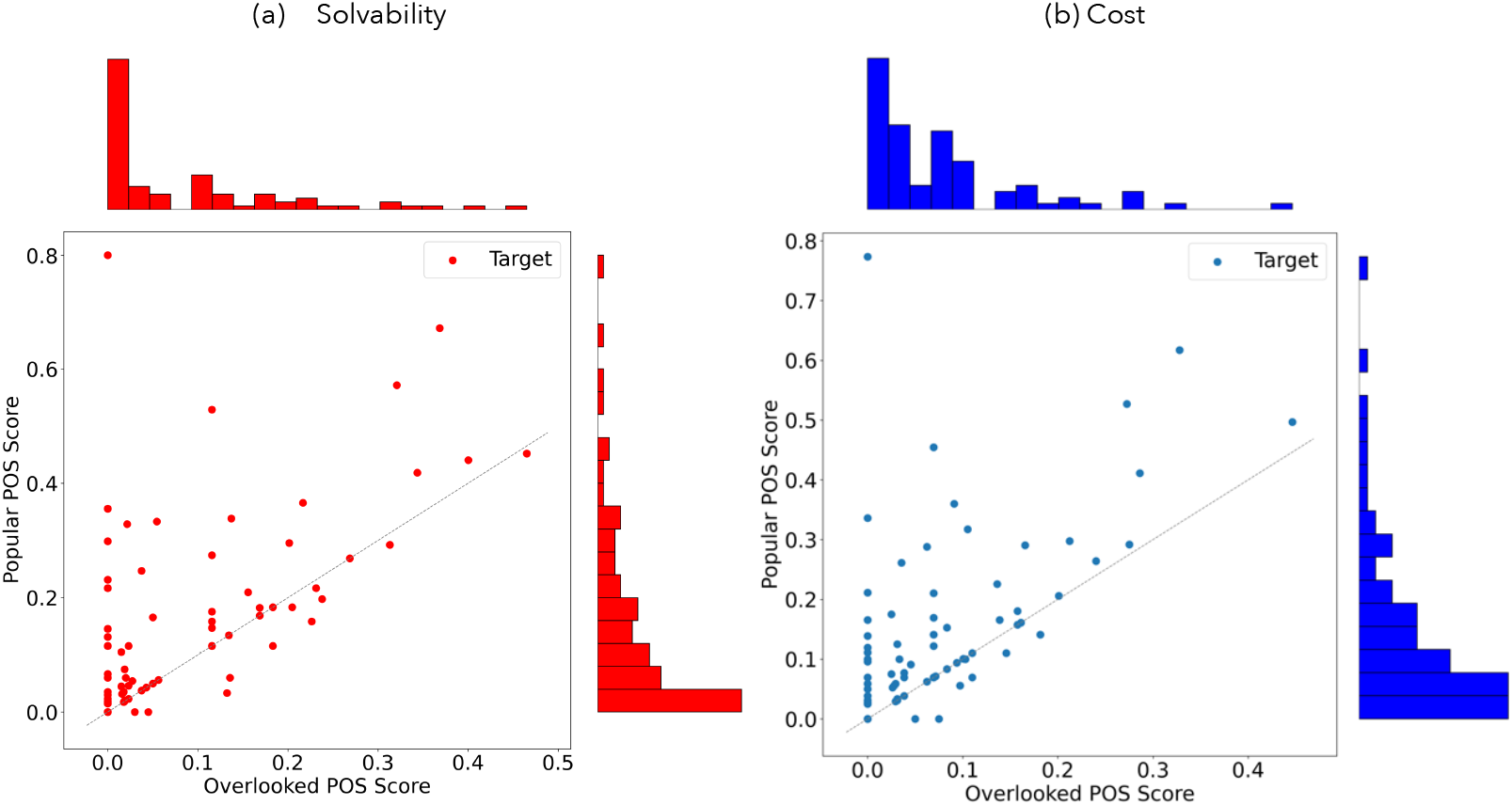
Impact of transformation optimisation on solvability and cost. Post-optimisation POS scores showing the effect of popular and overlooked transformation optimisation on solvability (a) and cost (b) when synthesising kinase target actives (each point is a target). In general, optimising popular transformations more often leads to improved performance, however, when overlooked templates are found, their promotion can lead to large increases in the number of molecules solved or reduction in synthesis cost. For some targets the same effect is seen after Popular or Overlooked template optimisation.

Similarly, when the goal is to reduce synthesis costs optimising Popular templates is often the most effective approach (Figure 3b). The reduced impact of Overlooked template optimisation suggests that promoting Overlooked templates may be less effective or that fewer unexplored strategies are available for certain target actives. Our findings support the latter, as there are often fewer Overlooked templates compared to Popular templates for the same target. A template is considered Overlooked when it appears in the lowest-cost route generated by the template-free method, and that route is less expensive than any template-based alternative.

However, this scenario is unlikely to occur for all targets actives. According to our criteria (see methods), we identified overlooked templates for 83% of molecule sets, whereas popular templates are identified for all molecule sets. Despite this, we still observed instances where Overlooked template optimisation showed positive effects, even where Popular template optimisation was ineffective. This result indicates that while it may be more challenging for the retrosynthesis model to incorporate Overlooked templates effectively, their promotion and inclusion can have a significant impact on synthesis performance. Our results are target-specific highlighting the importance of tailoring searches to necessary chemical space. Figure 3 also shows a moderate correlation between Overlooked and Popular template optimisation. The Pearson correlation coefficients for solvability (*r*(78) = 0.587, p < 0.001) and cost (*r*(78) = 0.597, p < 0.001) support this observation. These correlations suggest that Overlooked and Popular templates often represent similar chemical transformations between reagents and products, which may lead to analogous outcomes in route search performance.

We utilised retrosynthesis tools to identify under-utilised reactions that, when prioritised, can lead to improved synthesis pathways. These results demonstrate the potential for combining current retrosynthesis tools to more effectively explore and utilise reaction space, ultimately enabling the synthesis of more molecules or reducing costs.

### Integrating novel chemistry

So far, we have focused on promoting already documented reactions without introducing novel chemistry. Template-free tools are not restricted to selecting documented templates during route searches they can generate transformations regressively based on training data. This approach can result in the identification of Popular templates, Overlooked templates present in reaction libraries, and Novel templates that feature new chemistry that may not yet be feasible (Table 2).

To explore the potential benefits of integrating new chemistry, we conducted an experiment optimising novel templates when synthesising 20 active molecules for 80 kinase targets (see methods for details). We then repeated route searches promoting these new templates and compared the results to standard retrosynthesis. At least one novel template was identified for all 80 kinase targets. Upon optimisation of these novel templates, 66 of the 80 targets yielded new routes incorporating the novel templates, with better route performance than the best pre-optimisation route. Highlighting, potentially useful alternative synthesis pathways.

Furthermore, the routes that incorporated a novel template, lead to an average ∼ 26% reduction in cost.

To showcase the versatility of our method, in the following work we employed a metric, defined by Equation 2, to determine the cost of a synthesis pathway. This metric considers the number of steps, and the precursors required, offering a simple and rapid calculation that is independent of precursor stock library.

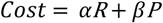

*Equation 2: Cost function used to determine the cost of a synthesis pathway in novel template optimisation experiment. R is the number of reaction steps, and P is the number of purchasable precursors. In our experiment, α = 0.7 and β = 0.3.*

As a case study, Figure 4a illustrates the cheapest route to a target molecule before template optimisation. In this reaction, five precursors are used in a five-step processes: first an intramolecular cyclisation reaction between the ethyl ester and pyrimidine-amine results in the formation of a bicyclic lactam followed by sulphoxide formation using an oxaziridine oxidising agent. At the same time an amide formation reaction occurs between an acyl chloride and amine, before reacting with the sulphoxide in an aromatic nucleophilic substitution. Finally, the secondary amine undergoes a deprotection step to remove a carboxybenzyl group. Alternatively, the optimised pathway (Figure 4b) is a much simpler four-step synthesis with only three purchasable precursors. In this route, two novel templates are utilised (orange boxes), first the direct oxidation the methylthio group, followed by the selective substitution of the sulphoxide group without the need of a protecting group on the terminal secondary amine. The reduction in the number of steps results in a route that is 40% cheaper using our cost function (described above and in methods, performance evaluation). In addition, if we calculate the route price from Molport, the optimised route is 25% cheaper than the original.

**Figure 4.**
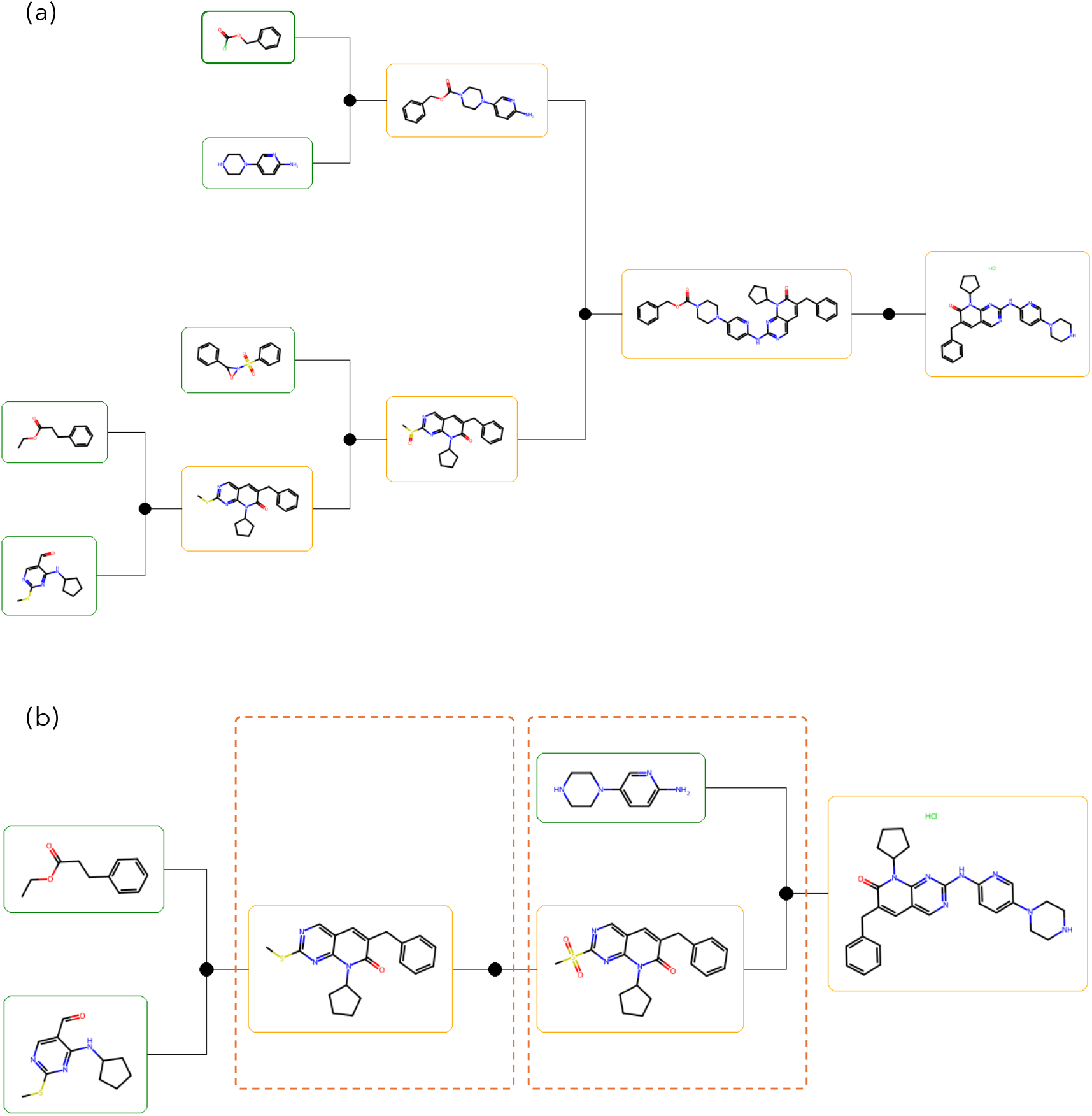
Effect of inclusion of novel templates into synthesis planning. Synthesis pathways to the same target active, (a) before novel template optimisation and (b) after optimisation. The optimised route is more efficient with fewer steps and purchasable precursors. Additionally, the cost of precursors from Molport is 25% cheaper than the original route.

This novel route is not without potential challenges, such as selectivity and reactivity issues, nonetheless, it represents a data driven approach to establishing the chemical reactions needed to reduce compound synthesis costs. Our template identification method highlighted this specific transformation as the most crucial to develop, as it proved most effective in reducing synthesis costs within the constraints of the precursor library and other available transformations. These findings demonstrate the significant potential of integrating novel chemistry into retrosynthesis tools. By expanding beyond documented reactions, we can unlock new, more efficient pathways to target molecules, particularly in less explored areas of chemical space. This approach not only enhances the capabilities of existing retrosynthesis tools but also points towards promising areas for future reaction development in synthetic organic chemistry.

## Conclusion

Our investigation has highlighted the challenge that current retrosynthesis tools face in efficiently exploring reaction space, particularly for novel or complex molecules in less explored areas of chemical space. Our results show that current tools tend to fail when reactions from under-utilised areas of reaction space are necessary for success. The correlation between retrosynthesis performance and the familiarity of chemical space underscores the need for more creative tools that can extend beyond standard synthesis practices.

Our method for integrating novel and overlooked synthesis strategies and reactions showed that retrosynthesis performance depends on several factors, including reaction diversity, molecule type, and synthesis goals. These results also illustrate the benefits of combining template-free approaches with template-based methods to expand the scope and efficiency of synthesis planning, emulating the actions of a more creative expert chemist. In addition to solving routes to target molecules, we have shown that our tool can highlight the chemical reactions that should be developed to improve our navigation of chemical space.

## Methods

### Datasets

We used ChEMBL^48^ as our source of target active molecules. We searched their database for known actives against kinase and non-kinase targets. We randomly collected ChEMBL IDs for kinase and non-kinase targets that have at least 50 known actives. Then randomly selected 50 molecules for each target. This formed the basis of the kinase and non-kinase collection of molecules for our comparison of template-based and template-free methods. From these datasets 750 molecules were randomly selected for processing with Postera and AiZynthFinder (see S5 for more details).

In our integration of template-based and template-free methods, we used a smaller dataset of molecules. For our popular and overlooked strategy experiments we randomly selected 80 kinase and non-kinase targets from ChEMBL, then collected 10 actives for each target. For our integration of novel chemistry, we randomly selected 20 kinase targets from ChEMBL, then randomly selected 20 actives molecules for each target for retrosynthesis.

### Property characterization

Molecular weight (MW), topological polar surface (TPSA) and QED were calculated using the chemical descriptor module from RDKit.^49^ Synthetic accessibility was calculated using SAscore.^50^ Molecular Tanimoto similarities were calculated using RDKit.^49^

### Retrosynthesis

We assessed the performance of two retrosynthesis tools. The first was the template-based AiZynthFinder^51^, the second was the template-free Postera.^30^ The aim of these tools is to breakdown a target molecule into a set of predefined purchasable precursors (e.g. zinc^52^). They do this by identifying a series of reactions (or transformations) that if applied in reverse break the molecule iteratively till a precursor is generated. These reactions are captured as templates, represented by SMARTS.^53^

AiZynthFinder is an open-source retrosynthesis tools that utilises a pathfinding algorithm (e.g. Monte-Carlo tree search or A*) with a neural expansion network to find synthetic routes for target molecules. The central template-based algorithm uses a neural network to guide the tree-search. At each iteration, the most promising molecule (node) is chosen, then the network uses a policy to suggest the best transformation to apply to the molecule. This process is repeated until a maximum depth is reached, limit elapses or a purchasable precursor is found. Full details of the model and networks used are available in.^51,54^

Postera is a template-free retrosynthesis tools based on the molecular transformer.^31,55^ We used their v1 retrosynthesis endpoint with a maximum search depth of four and the Molport^56^ building block library. For AiZynthFinder we implemented a maximum tree search depth of four to match Postera. All other parameters were kept as default. For consistency and to allow reasonable comparison to Postera, the Molport building block catalog was also used.

### Performance evaluation

In addition to the POS score described in the results, we used three simple metrics to evaluate retrosynthesis performance:

1. Solvability: the total number of solved molecules
2. Route Price: the sum of purchasable precursors, number of steps and precursors
3. Route Efficiency: combination of precursors and reaction steps

A route was solved if all precursors (terminal nodes) were purchasable from a stock library. These metrics were chosen as they were easily interpretable and computationally straightforward to calculate.

The POS scoring function is sensitive to different *α, β, and γ* values, therefore, the function can be adjusted to the users’ preferences. In our tool (see Code Availability), alternative metrics not discussed here can be used, for example precursor price can be determined using other tools, such as CoPriNet.^57^ Other metrics could in future be used to calculate cost - for example from recent work from Fromer et al.^58^

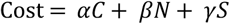

*Equation 3: Mathematical description of the route price. C is the total sum of precursor prices; N is the number of precursors and S is the number of reactions. For all our calculation a = 0*.*7, b = 0*.*15 and y = 0*.*15*.

Equation 2 shows how we calculated the route price as a combination precursor price, route length and number of precursors. The precursor price was the price range of 1mg sourced from the Molport library. Table 3 details how documented price ranges were translated to quantative values. We also explored the effect of search constraints on retrosynthesis performance detailed in S8.

**Table 3:**
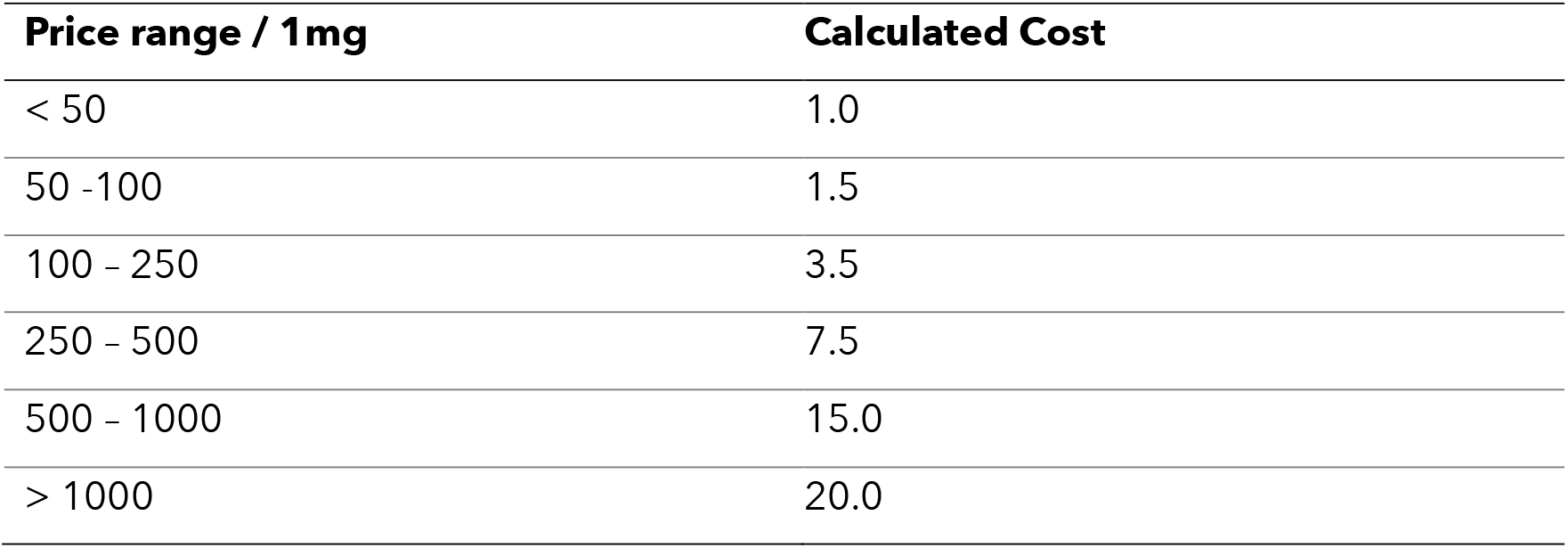
Conversion of Molport price ranges to discrete values for route cost calculations.

In addition to solvability and route price, we also used route efficiency in our novel template integration experiments (Equation 2). This metric allows us to evaluate the efficiency of synthesis pathways. This metric incorporates the number of reaction steps (R) and the number of purchasable precursors (P), enabling quick and independent assessment without reliance on a specific precursor stock library. By emphasising reaction efficiency, this approach provides valuable insights into how we could design more streamlined and cost-effective synthetic pathways.

## Template Analysis

### Representing Reactions and Template Similarity

In our study, reaction templates were represented as SMARTS. In template-based searches they were collected from the USPTO library available through AiynthFinder.^37^ In template-free searches, no template was provided, instead transformations were described as product and reaction SMILES pairs. To convert these SMILES to SMARTS for comparison, we used RDChiral.^59^ This generated templates with a radius of one which matches the template preparation used to generate the USPTO template library. To compare templates, we canonicalised extracted the templates using RDCanon^60^, then checked for matches with canonical forms of the USPTO templates (also generated using RDCanon). Templates were considered the same if their canonical forms were identical.

### Chemical Space Coverage

To display the diversity of reactions used by AiZynthFinder in solved and unsolved routes, we plotted the number of template clusters used at each step synthesis routes for kinase actives. We generated fingerprints for all templates using RDKit^49^, then clustered them using DBSCAN.^47^ The DBSCAN epsilon was 0.3, while the minimum samples per cluster was one. Epsilon defines the maximum distance between two samples for them to be neighbours, while the minimum sample parameter is the number of neighbouring samples needed for a point to be a core point of a cluster. These parameters were chosen after a series of experiments which aimed at getting meaningful clusters whilst reducing the number of noise points.

### Library Occurrence

We attempted to solve routes for 2,500 kinase active molecules. We collected all the successful routes for the 1666 solved kinase (67 % of the 2,500 attempted). AiZynthFinder used approximately 90,000 templates across all solved routes for all molecules, 4,905 of which were unique. We then collected all unique templates based on their frequency. Template frequency was then binned and the average USPTO occurrence for each bin was determined.

### Template Classification

We categorised reactions into four distinct reactions classes based on their use and availability in template-based and template-free searches on the same set of molecules (see S10 for diagram). Popular reactions are ranked based on how often they are used in the cheapest routes generated by template-based methods; these are well-documented and established reactions. Overlooked reactions are those available in the template-based reaction library but not used in a template-based search, however, they are used in the template-free search. These reactions highlight under-utilised synthetic strategies using documented reactions. Novel reactions are those that are absent from the template-based reaction libraries but are generated and successfully applied by template-free approaches, representing undocumented or entirely new chemistry. Novel templates can also include templates used in routes to molecules that were only solved by template-based methods. This reaction classification allows us to evaluate the contributions of different reaction types when integrating template-based and template-free approaches.

### Reaction Optimisation

We used AiZynthFinder to determine the effect of promoting the use of specific transformations during retrosynthesis. During the expansion stage of tree-search a neural network is trained on USPTO reaction data to suggest the best transformation to apply to the molecule at a node. This policy network calculates the likelihood of all templates giving each one a score between 0 and 1. More suitable transformations are given a higher score; however, this is biased by how often a transformation is used in reaction data. The prevalence of a reaction in the training data corresponds to how available or easy it is to use in practice because the more accessible a reaction is, the more often it will appear in reaction databases.

Our aim for optimisation was to simulate a reaction being more accessible in practice and to encourage the model to use specific transformations. To do this, we artificially increased the policy value for specific templates. This is analogous to optimising or improving the success of a reaction in reaction data. We did this in two ways. For popular and overlooked template optimisation, we determined the value of the highest likelihood transformation from the expansion model, then increased the policy value of optimised transformations to this value before normalizing the new policy values. This maintains the relative likelihoods of all other reactions. Alternatively, for novel template optimisation we calculated the new prior, *nP*, by increasing the original prior, *oP*, by an optimisation value, OV:

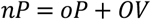

OV is between 0 and 1. It is calculated using the number of times a template is used in TF routes that are cheaper than TB routes (C), the total number of times it’s used (T), and the policy value range from the expansion model (R):

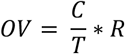

For novel templates, where an original prior is unavailable, we estimate a prior (eP) based on the minimum prior (mP), the usage ratio between cheap and total routes, and the range of original priors (R) from the expansion model.

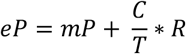

This estimation is proportional to how often a template is useful in cheap routes and will sit within the range of priors from the model. Thus, the new prior for novel templates is given by:

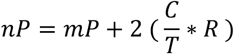

The original and integrated priors are then normalised to maintain appropriate limits. This method allows us to optimise all relevant templates relative to how useful they are predicted to be (optimisation factor). Neither of the methods used forces the model to select the optimised transformation; instead, it increases the likelihood of its selection.

### Popular vs Overlooked Strategies

We examined the effect of incorporating popular or overlooked synthesis strategies on retrosynthesis performance. A synthesis strategy is characterised by the most frequently used templates applied in search results. We selected 80 kinase and non-kinase targets, each with a minimum of 10 active molecules. We ran both our template-based and template-free methods and compared the results to identify popular and overlooked templates.

As popular and overlooked templates must be present in the TB template library, no undocumented templates were introduced. All identified templates were ranked based on their optimisation value. This is a ratio between the number of times each template was used in cheap routes and its’ overall occurrence (see methods: reaction optimisation). These templates were then optimised in a route search using AiZynthFinder.

### Integration of novel chemistry

To integrate novel strategies into template-based retrosynthesis, we expanded AiZynthFinder to include entirely new templates representing novel or undocumented chemistry not available in the USPTO reaction library. Again, we compared the template-based and template-free search results on our 80 targets (each with 20 active molecules) and identified popular, overlooked and novel reactions for optimisation.

This classification of popular, overlooked, and novel reactions enabled us to evaluate the contributions of different reaction types when integrating TB and TF approaches in retrosynthesis. The five most frequently used templates were selected and optimised for each target by adding them to the list of possible actions during expansion. Each optimised template was given the same policy value as the highest scoring template from the expansion model, increasing the likelihood of its selection without forcing its use. Route searches for all 20 active molecules from each kinase and non-kinase target were completed before and after optimisation of each reaction class.

## Code availability

The tool and code used in this study is available at https://github.com/m-mokaya/retromix.git

## Acknowledgements

This work was supported by funding from the Engineering and Physical Sciences Research Council (EPSRC) and Exscientia.

## References

1. Rennane, S., Baker, L. & Mulcahy, A. Estimating the Cost of Industry Investment in Drug Research and Development: A Review of Methods and Results. INQUIRY 58, 00469580211059731 (2021).

2. Simoens, S. & Huys, I. R&D Costs of New Medicines: A Landscape Analysis. Frontiers in Medicine 8, (2021).

3. Brown, D. G., Wobst, H. J., Kapoor, A., Kenna, L. A. & Southall, N. Clinical development times for innovative drugs. Nature Reviews Drug Discovery 21, 793–794 (2021).

4. Romano, J. D. & Tatonetti, N. P. Informatics and Computational Methods in Natural Product Drug Discovery: A Review and Perspectives. Front. Genet. 10, 368 (2019).

5. Lin, X., Li, X. & Lin, X. A Review on Applications of Computational Methods in Drug Screening and Design. Molecules 25, (2020).

6. Mokaya, M. et al. Testing the limits of SMILES-based de novo molecular generation with curriculum and deep reinforcement learning. Nat Mach Intell 5, 386–394 (2023).

7. Blaschke, T. et al. REINVENT 2.0: An AI Tool for De Novo Drug Design. J. Chem. Inf. Model. 60, 5918–5922 (2020).

8. Deng, J. et al. A systematic study of key elements underlying molecular property prediction. Nat Commun 14, 6395 (2023).

9. Pang, C., Qiao, J., Zeng, X., Zou, Q. & Wei, L. Deep Generative Models in De Novo Drug Molecule Generation. J. Chem. Inf. Model. 64, 2174–2194 (2024).

10. Gao, W. & Coley, C. W. The Synthesizability of Molecules Proposed by Generative Models. 2002.07007 [cs, q-bio, stat] (2020).

11. Coley, C. W., Green, W. H. & Jensen, K. F. Machine Learning in Computer-Aided Synthesis Planning. Acc. Chem. Res. 51, 1281–1289 (2018).

12. Jiang, Y. et al. Artificial Intelligence for Retrosynthesis Prediction. Engineering 25, 32–50 (2023).

13. Sun, Y. & Sahinidis, N. V. Computer-aided retrosynthetic design: fundamentals, tools, and outlook. Current Opinion in Chemical Engineering 35, 100721 (2022).

14. Corey, E. J. & Wipke, W. T. Computer-Assisted Design of Complex Organic Syntheses. Science 166, 178–192 (1969).

15. Corey, E. J., Wipke, W. T., Cramer, R. D. & Howe, W. J. Computer-assisted synthetic analysis. Facile man-machine communication of chemical structure by interactive computer graphics. J. Am. Chem. Soc. 94, 421–430 (1972).

16. Irwin, R., Dimitriadis, S., He, J. & Bjerrum, E. J. Chemformer: a pre-trained transformer for computational chemistry. Mach. Learn.: Sci. Technol. 3, 015022 (2022).

17. Chen, S. & Jung, Y. Deep Retrosynthetic Reaction Prediction using Local Reactivity and Global Attention. JACS Au 1, 1612–1620 (2021).

18. Tetko, I. V., Karpov, P., Van Deursen, R. & Godin, G. State-of-the-art augmented NLP transformer models for direct and single-step retrosynthesis. Nat Commun 11, 5575 (2020).

19. Somnath, V. R., Bunne, C., Coley, C., Krause, A. & Barzilay, R. Learning Graph Models for Retrosynthesis Prediction. in Advances in Neural Information Processing Systems vol. 34 9405–9415 (Curran Associates, Inc., 2021).

20. Shi, C., Xu, M., Guo, H., Zhang, M. & Tang, J. A Graph to Graphs Framework for Retrosynthesis Prediction. in Proceedings of the 37th International Conference on Machine Learning 8818–8827 (PMLR, 2020).

21. Wang, X. et al. RetroPrime: A Diverse, plausible and Transformer-based method for Single-Step retrosynthesis predictions. Chemical Engineering Journal 420, 129845 (2021).

22. Kocsis, L. & Szepesvári, C. Bandit Based Monte-Carlo Planning. in Machine Learning: ECML 2006 (eds. Fürnkranz, J., Scheffer, T. & Spiliopoulou, M.) 282–293 (Springer, Berlin, Heidelberg, 2006). doi:10.1007/11871842_29.

23. Coulom, R. Efficient Selectivity and Backup Operators in Monte-Carlo Tree Search. in Computers and Games (eds. Van Den Herik, H. J., Ciancarini, P. & Donkers, H.H.L.M.) vol. 4630 72–83 (Springer Berlin Heidelberg, Berlin, Heidelberg, 2007).

24. Hart, P., Nilsson, N. & Raphael, B. A Formal Basis for the Heuristic Determination of Minimum Cost Paths. IEEE Trans. Syst. Sci. Cyber. 4, 100–107 (1968).

25. Segler, M. H. S., Preuss, M. & Waller, M. P. Planning chemical syntheses with deep neural networks and symbolic AI. Nature 555, 604–610 (2018).

26. Karpov, P., Godin, G. & Tetko, I. V. A Transformer Model for Retrosynthesis. in Artificial Neural Networks and Machine Learning – ICANN 2019: Workshop and Special Sessions (eds. Tetko, V., Kůrková, V., Karpov, P. & Theis, F.) 817–830 (Springer International Publishing, Cham, 2019). doi:10.1007/978-3-030-30493-5_78.

27. Coley, C. W. et al. A robotic platform for flow synthesis of organic compounds informed by AI planning. Science 365, eaax1566 (2019).

28. Kreutter, D. & Reymond, J.-L. Multistep retrosynthesis combining a disconnection aware triple transformer loop with a route penalty score guided tree search. Chem. Sci. 14, 9959– 9969 (2023).

29. Chen, B., Li, C., Dai, H. & Song, L. Retro*: Learning Retrosynthetic Planning with Neural Guided A* Search. 2006.15820 [cs, stat] (2020).

30. PostEra, Medicinal Chemistry Powered by Machine Learning. https://app.postera.ai/.

31. Schwaller, P. et al. Molecular Transformer: A Model for Uncertainty-Calibrated Chemical Reaction Prediction. ACS Cent. Sci. 5, 1572–1583 (2019).

32. Toniato, A. et al. Quantum chemical data generation as fill-in for reliability enhancement of machine-learning reaction and retrosynthesis planning. Digital Discovery (2023) doi:10.1039/D3DD00006K.

33. Sumiya, Y., Harabuchi, Y., Nagata, Y. & Maeda, S. Quantum Chemical Calculations to Trace Back Reaction Paths for the Prediction of Reactants. JACS Au 2, 1181–1188 (2022).

34. Ree, N., Göller, A. H. & Jensen, J. H. Automated quantum chemistry for estimating nucleophilicity and electrophilicity with applications to retrosynthesis and covalent inhibitors. Digital Discovery 3, 347–354 (2024).

35. Kadam, J.J. Quantum Machine Learning Technique for Automatic Retrosynthetic Reaction Pathway Search Method. International Journal on Recent and Innovation Trends in Computing and Communication 11, 2111–2122 (2023).

36. Liu, G. et al. Retrosynthetic Planning with Dual Value Networks. in Proceedings of the 40th International Conference on Machine Learning 22266–22276 (PMLR, 2023).

37. Guo, J. et al. Retrosynthesis Zero: Self-Improving Global Synthesis Planning Using Reinforcement Learning. J. Chem. Theory Comput. 20, 4921–4938 (2024).

38. Kearnes, S. M. et al. The Open Reaction Database. J. Am. Chem. Soc. 143, 18820–18826 (2021).

39. Zhang, C. & A. Lapkin, A. Reinforcement learning optimization of reaction routes on the basis of large, hybrid organic chemistry–synthetic biological, reaction network data. Reaction Chemistry & Engineering 8, 2491–2504 (2023).

40. Genheden, S. & Bjerrum, E. PaRoutes: towards a framework for benchmarking retrosynthesis route predictions. Digital Discovery 1, 527–539 (2022).

41. Zhong, Z. et al. Recent advances in deep learning for retrosynthesis. WIREs Computational Molecular Science n/a, e1694.

42. Roughley, S. D. & Jordan, A. M. The medicinal chemist’s toolbox: an analysis of reactions used in the pursuit of drug candidates. J Med Chem 54, 3451–3479 (2011).

43. Schwaller, P., Petraglia, R., Nair, V. H. & Laino, T. Evaluation Metrics for Single-Step Retrosynthetic Models.

44. Santos, R. et al. A comprehensive map of molecular drug targets. Nat Rev Drug Discov 16, 19–34 (2017).

45. D, L. Chemical Reactions from US Patents (1976–Sep 2016). (2017).

46. Lowe, D. M. Extraction of chemical structures and reactions from the literature. (University of Cambridge, 2012).

47. Ester, M., Kriegel, H.-P., Sander, J. & Xu, X. A Density-Based Algorithm for Discovering Clusters in Large Spatial Databases with Noise.

48. Gaulton, A. et al. ChEMBL: a large-scale bioactivity database for drug discovery. Nucleic Acids Res 40, D1100–D1107 (2012).

49. Landrum, G. RDKit: Open-source cheminformatics. (2006).

50. Ertl, P. & Schuffenhauer, A. Estimation of synthetic accessibility score of drug-like molecules based on molecular complexity and fragment contributions. Journal of Cheminformatics 1, 8 (2009).

51. Genheden, S. et al. AiZynthFinder: a fast, robust and flexible open-source software for retrosynthetic planning. J Cheminform 12, 70 (2020).

52. Irwin, J. J. et al. ZINC20—A Free Ultralarge-Scale Chemical Database for Ligand Discovery. J. Chem. Inf. Model. 60, 6065–6073 (2020).

53. Daylight Theory: SMARTS - A Language for Describing Molecular Patterns. https://www.daylight.com/dayhtml/doc/theory/theory.smarts.html.

54. Segler, M. H. S. & Waller, M. P. Neural-Symbolic Machine Learning for Retrosynthesis and Reaction Prediction. Chemistry – A European Journal 23, 5966–5971 (2017).

55. Lee, A. A. et al. Molecular Transformer unifies reaction prediction and retrosynthesis across pharma chemical space. Chem. Commun. 55, 12152–12155 (2019).

56. Molport. Molport Molecular Database.

57. Sanchez-Garcia, R. et al. CoPriNet: graph neural networks provide accurate and rapid compound price prediction for molecule prioritisation. Digital Discovery 2, 103–111 (2023).

58. Fromer, J. C. & Coley, C. W. An algorithmic framework for synthetic cost-aware decision making in molecular design. Nat Comput Sci 4, 440–450 (2024).

59. Coley, C. W., Green, W. H. & Jensen, K. F. RDChiral: An RDKit Wrapper for Handling Stereochemistry in Retrosynthetic Template Extraction and Application. J. Chem. Inf. Model. 59, 2529–2537 (2019).

60. Mahjour, B. A. & Coley, C. W. RDCanon: A Python Package for Canonicalizing the Order of Tokens in SMARTS Queries. J. Chem. Inf. Model. 64, 2948–2954 (2024).

