## Supplementary Information for "Expanding the Synthome: Generating and integrating novel reactions into retrosynthesis tools"

#### S1 Molecular Properties of Solved and Unsolved Molecules

It could be argued that kinase actives are easier to synthesise than their non-kinase counterparts. We do not believe this to be entirely true. In Table 1 we present the mean values of three molecular properties (molecular weight, QED and TPSA). The averages of solved molecules were similar across active sets (e.g. 385 Da and 391 Da respectively for molecular weight). We do see unfavourable shifts in these values in unsolved molecules for both kinase and non-kinase actives. These shifts are often larger for non-kinase unsolved molecules and will contribute to their synthetic accessibility. However, Figure 1a shows there is a substantial overlap in the property distributions of solved and unsolved molecules. For example, most unsolved molecules have a molecular weight within the molecular weight range of solved molecules (ca. 100 – 750 Da). We do not observe a large general difference in the properties of solved and unsolved molecules. This implies that while some molecules may have characteristics that are harder to synthesize, these differences alone are not the cause of retrosynthetic failure. For our assessment of the limitations of retrosynthesis tools to hold, both kinase and non-kinase actives must be broadly accessible.

[1] Department of Statistics, University of Oxford

[2] Department of Chemistry, University of Liverpool

[3] Department of Computer Science, University of Liverpool

*Table 1: Molecular properties of kinase and non-kinase actives. In general, similar values are observed for kinase and non-kinase actives. Non-kinase actives do exhibit properties that are harder to synthesis, however, these are not enough to explain poorer retrosynthesis performance.*

| **Target Class** | **Property** | | | | | |
|---|---|---|---|---|---|---|
| | **TPSA** | | **MW** | | **QED** | |
| | Solved | Unsolved | Solved | Unsolved | Solved | Unsolved |
| Kinase | 82.8 | 121 | 385 | 457 | 0.576 | 0.467 |
| Non-Kinase | 83.9 | 211 | 391 | 709 | 0.521 | 0.373 |

Importantly, we do see a shift in the synthetic accessibility (SA) score of solved and unsolved molecules. Figure 1b, shows that there are significantly more molecules that with lower synthetic accessibility (higher numerical value – see methods), that are unsolved using AiZynthFinder. This evidence is consolidated by the average SA score for unsolved and solved molecules. For kinases it is 2.66 solved and 3.30 for unsolved. For

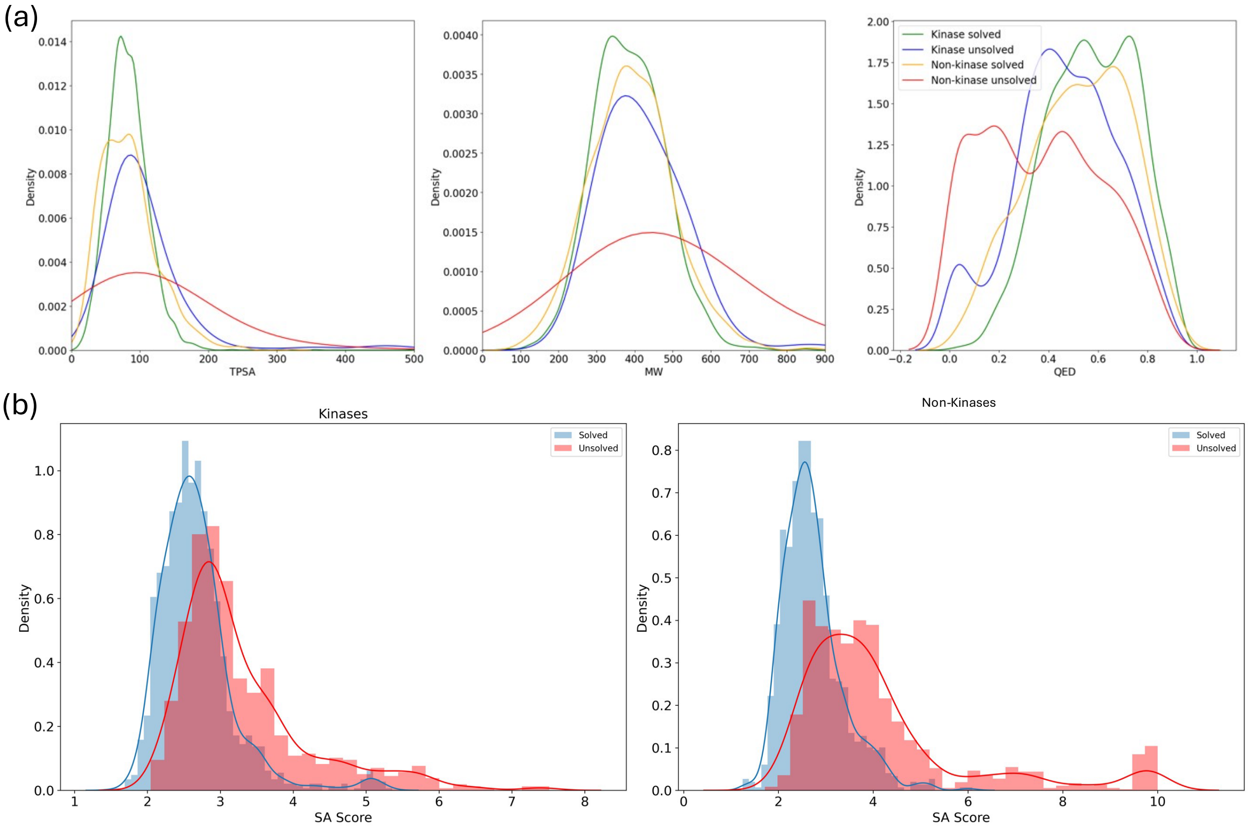


*Figure 1: Molecular property distributions of solved and unsolved molecules. (a) TPSA, Molecular Weight (MW) and QED distributions for actives against kinase and non-kinase targets. (b) Distribution of SA score for actives against kinase and non-kinase targets.*

all non-kinase activities, it is 2.71 for solved and 4.14 for unsolved. The observed shift aligns with the anticipated relationship between SA score and fragment complexity, where a molecules' SA score is inversely proportional to the presence of its fragments in chemical libraries.[2] Better documented moieties are perceived as less complex, leading to lower SA scores. Therefore, the observed reduction in synthetic accessibility for less well researched, thus harder-to-solve molecules follows as expected.[3]

#### S2 AiZynthFinder template-based vs template-free

AiZynthFinder is an open-source retrosynthesis tool that is available as a template-based and template-free tool.[1] Our work is based on the template-based variant. We compare its performance with another template-free tool, Postera's Manifold.[4] We find that the template-based AiZynthFinder does not perform as well as the template-free variant on kinase and non-kinase target actives. We posit that this is, in part, due to limitations that the template-based tools have. To verify this, we compared the template-based and template-free AiZynthFinder variants on 300 kinase and non-kinase molecules.

*Table 2: Performance of template-based and template-free AiZynthFinder on a set of kinase and non-kinase actives. Performance for both tools was better on kinase targets relative to non-kinase targets. Both and neither shows the percentage of molecules attempted by both methods that had solved or unsolved routes.*

| **Target Class** | **Percentage Solved / %** | | | |
|---|---|---|---|---|
| | **AiZ – TB** | **AiZ – TF** | **Both** | **Neither** |
| Kinase | 69.0 | 72.3 | 51.0 | 11.0 |
| Non-kinase | 53.3 | 64.0 | 46.7 | 30.7 |

In line with our main results, we find that the template-free tool performs better than the template-based alternative. We also found that both methods perform better on kinase target actives compared to non-kinase target actives. This further supports our hypothesis that the tools tested perform better on well explored areas of chemical space, however they struggle on less well explored areas.

#### S3 Retrosynthesis failure modes

Understanding the cause of failure is important when trying to understand the limitations of retrosynthetic tools. Our results show that these failures are likely attributed to the models' inability to successfully use less popular transformations, preventing the resolution of intermediates to purchasable building blocks. To test whether this is true, we collected all the failed routes for kinase and non-kinase target actives using AiZynthFinder.[1] For each route we categorised its reason for failure such as search time and MCTS (Monte Carlo Tree Search) tree search depth. For this experiment we used AiZynthFinder with default search parameters.

We determined four failure modes:

1. No routes were found while within set limits (None).
2. No routes were found because the search time limit was reached (Time).
3. No routes were found because the maximum search tree depth was reached (Depth).
4. No routes were found because both limits were reached (Both).

Of the 750 of kinases and non-kinase actives tested, only one molecule was unresolved due to a search time limitation (Table 3). Therefore, inappropriate building blocks or templates were ultimately the causes of failure. This could be attributed to the expansion model's limitations in recommending suitable templates for generated intermediates or the absence of templates applicable to non-purchasable terminal nodes within default search parameters. This may be due to a combination of factors. Firstly, the expansion model is unable to identify appropriate templates to apply to intermediates. Secondly, the templates needed to resolve routes given the set of available building blocks do not exist. We suggest that a combination of both is likely.

*Table 3: Reasons for retrosynthesis failure. Highlights how inappropriate templates or building blocks are often the primary causes. 'None' indicates an incomplete synthesis pathway while the parameters of the search tree are within set limits (1). 'Time' signifies an incomplete synthesis pathway due to reaching the search time limit (2). 'Depth' indicates an incomplete synthesis pathway because the maximum search tree depth was reached (3). 'Both' implies an incomplete synthesis pathway due to reaching both the search depth and time limits (4).*

| **Target Class** | **Reason for Failure / %** |
|---|---|

| | None | Time | Depth | Both |
|---|---|---|---|---|
| Kinase | 50.1 | 0.00 | 49.6 | 0.240 |
| Non – kinase | 52.3 | 0.133 | 47.7 | 0.00 |

#### S4 Template Use and Library Occurrences

We compared the library occurrences within the USPTO reaction database of templates used to solve 750 kinase and non-kinase molecules. In our paper we show that on average, the more a template is used, the more often that template appears in the reaction library. Here, we show that for unsolved routes, we see more templates with lower template occurrences used in unsolved routes, especially for non-kinase targets. This suggests that less popular templates were used more often in routes that ultimately failed.

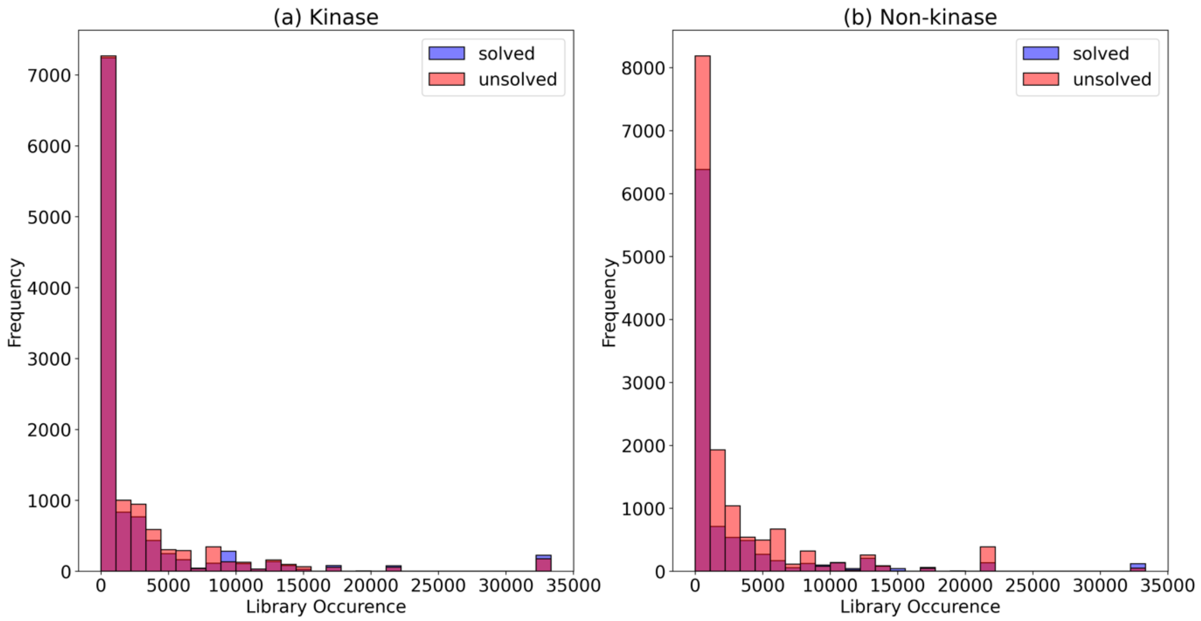


*Figure 2: Library occurrences distributions of templates in solved (blue) and unsolved (red) routes for (a) kinase and (b) non-kinase target actives.*

#### S5 Target and Active Data

All kinase and non-kinase targets along with associated actives are available in the 'data' section of our repository: https://github.com/m-mokaya/retromix.git

#### S6 Alternative POS weights

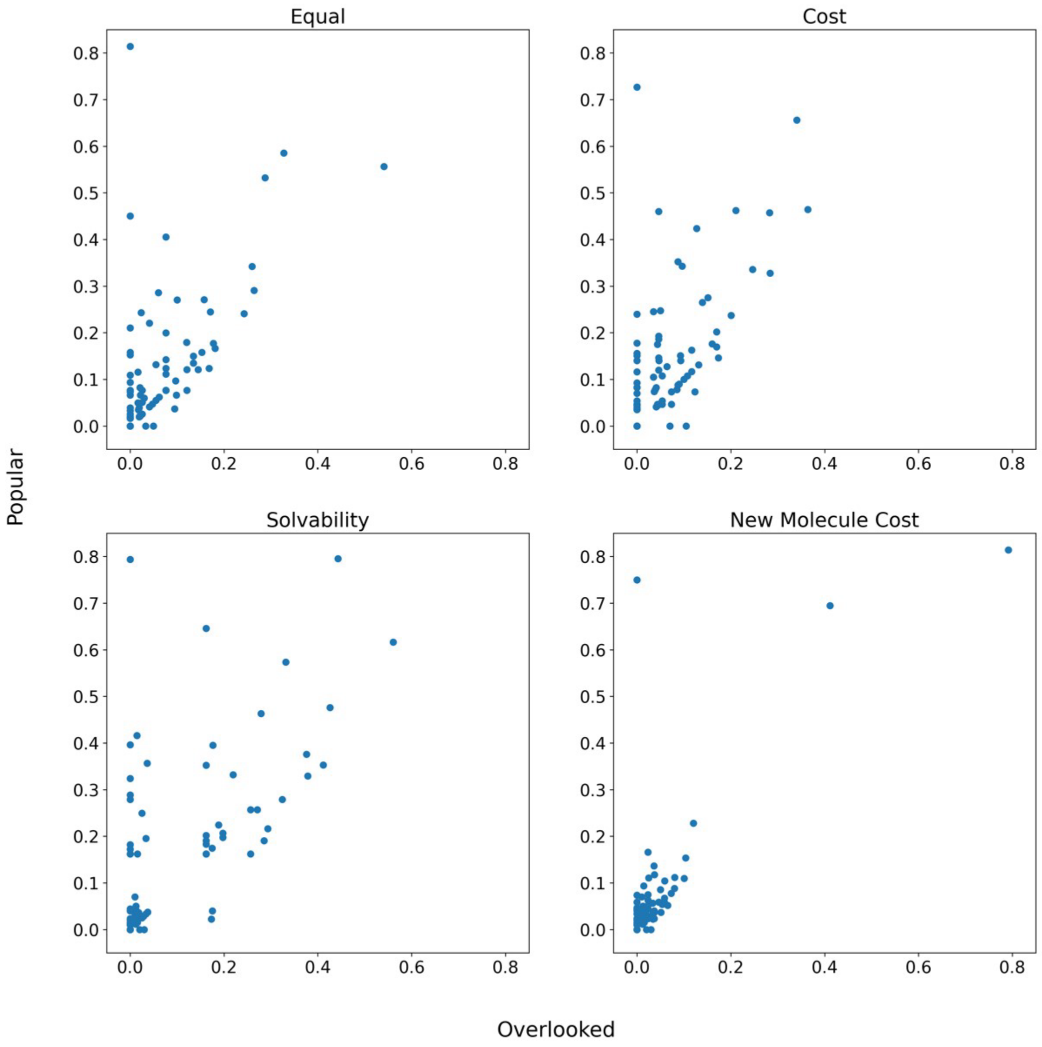


*Figure 3 : POS scores for kinase targets calculated using different POS coefficients . Each represents a different objective. (A) a, b and y are equal and 0.33, (b) cost prioritised POS scores. a = 0.7, b = 0.1 and y = 0. 2. (c) solvability prioritised POS scores. a = 0.2, b = 0.1, y = 0.7. (d) new molecule cost prioritised POS scores. a = 0.2, b = 0.7, y = 0.1.*

### S7 Comparing popular or novel strategies

We tried multiple template identification methods. In our paper, we optimise all popular and novel templates, however, their optimisation factor is dependent on how often they are used in the cheapest routes. Alternatively, we also tried to limit optimisation to the top 5 most used templates for each class. Here the most frequently used templates across all routes, independent of price, were optimised. In the expansion phase, their likelihood was increased to match the highest scoring template from the expansion model.

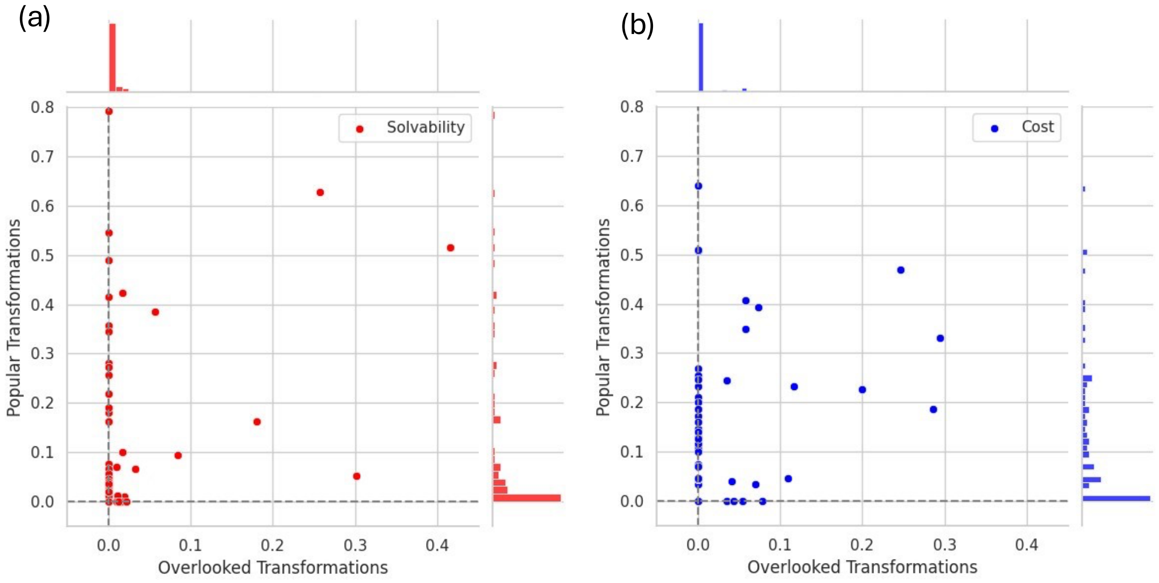


*Figure 4: Impact of transformation optimisation on solvability and cost. Post-optimisation POS scores showing the effect of popular and overlooked transformation optimisation on solvability (a) and cost (b) when synthesising kinase target actives. In general, optimi sing popular transformations more often leads to improved performance, however, when overlooked templates are found, their promotion can lead to large increases in the number of molecules solved or reduction in synthesis cost.*

In this setup, we see more targets were with zero POS scores for either template class. This is likely because, we are optimising fewer templates, and we have not identified any templates that will lead to positive effects. Therefore, we prefer to optimise more template with variable optimisation factors.

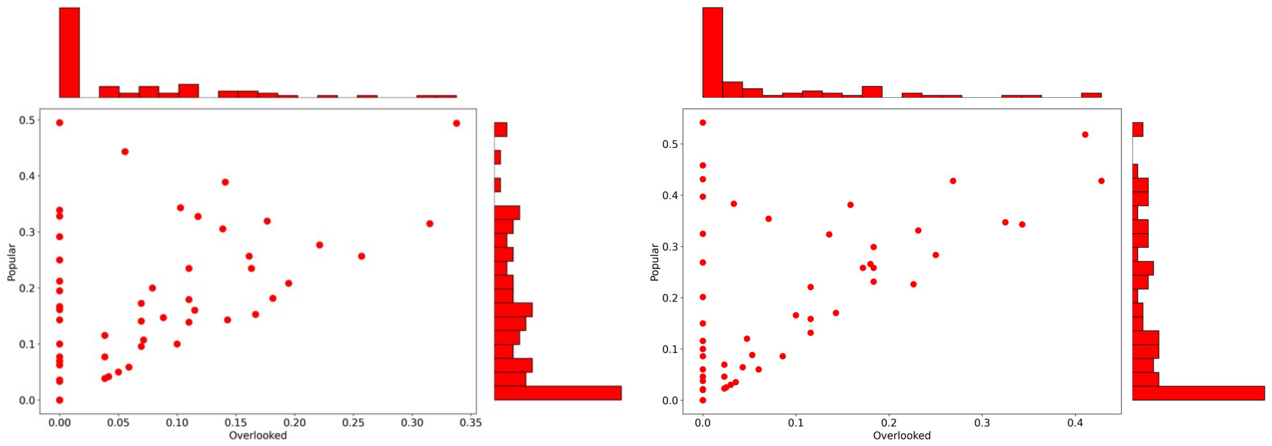


*Figure 5: Impact of transformation optimisation on solvability and cost. Post -optimisation POS scores showing the effect of popular and overlooked transformation optimisation on solvability (a) and cost (b) when synthesising non-kinase target actives.*

#### S8 Retrosynthesis failure modes

Understanding the cause of failure is important when trying to understand the limitations of retrosynthetic tools. Our results show that these failures are likely attributed to the models' inability to successfully use less popular transformations, preventing the resolution of intermediates to purchasable building blocks. To test whether this is true, we collected all the failed routes for kinase and non-kinase target actives using AiZynthFinder.[1] For each route we categorised its reason for failure such as search time and MCTS (Monte Carlo Tree Search) tree search depth. For this experiment we used AiZynthFinder with default search parameters.

We determined four failure modes:

1. No routes were found while within set limits (None).
2. No routes were found because the search time limit was reached (Time).
3. No routes were found because the maximum search tree depth was reached (Depth).
4. No routes were found because both limits were reached (Both).

Of the 750 of kinases and non-kinase actives tested, only one molecule was unresolved due to a search time limitation (Table 3). Therefore, inappropriate building blocks or templates were ultimately the causes of failure. This could be attributed to the expansion model's limitations in recommending suitable templates for generated intermediates or the absence of templates applicable to non-purchasable terminal nodes within default search parameters. This may be due to a combination of factors. Firstly, the expansion model is unable to identify appropriate templates to apply to intermediates. Secondly, the templates needed to resolve routes given the set of available building blocks do not exist. We suggest that a combination of both is likely.

*Table 4: Reasons for retrosynthesis failure. Highlights how inappropriate templates or building blocks are often the primary causes. 'None' indicates an incomplete synthesis pathway while the parameters of the search tree are within set limits (1). 'Time' signifies an incomplete synthesis pathway due to reaching the search time limit (2). 'Depth' indicates an incomplete synthesis pathway because the maximum search tree depth was reached (3). 'Both' implies an incomplete synthesis pathway due to reaching both the search depth and time limits (4).*

| **Target Class** | **Reason for Failure / %** | | | |
|---|---|---|---|---|
| | **None** | **Time** | **Depth** | **Both** |
| Kinase | 50.1 | 0.00 | 49.6 | 0.240 |
| Non – kinase | 52.3 | 0.133 | 47.7 | 0.00 |

#### S9 Standard vs Explore

We experimented with the search parameters for AiZynthFinder as it could be argued that any failures could be resolved by encouraging AiZynthFinder to explore more during the search process. If the tools' ability to pick appropriate templates is a limiting factor in performance, relaxing the search constraints should lead to improved performance. However, with no introduction of new templates or building blocks we still expect no routes to be found for some molecules.

*Table 5: Effects of expanding the search constraints for AiZynthFinder leads to more solved molecules but less favourable routes. Overall, AiZynthFinders' route solving performance increases, however, the route length and number of purchasable precursors for the molecules now solved (that weren't previously) is higher.*

| **Status** | **Percentage solved / %** | **Av. Top Route Length** | **Av. Num. Purchasable Blocks** |
|---|---|---|---|
| Standard | 66.6 | 2.04 | 3.00 |
| Explore | 83.2 | 3.40 | 4.03 |

Table 5 highlights how using more permissive parameters does lead to an improvement in route finding performance. Here we ran the same route search on kinase actives with relaxed search criteria. The number of transforms allowed per route was increased from 4 to 12, exploration coefficient was increased from 1.4 to 6.0, iteration limit was increased from 100 to 200 and the top 200 templates from the policy are considered (compared to 50 in the standard case). We report that under these relaxed criteria, there was an increase in the number of solved molecules. However, the top routes generated for newly solved molecules are on average 66% longer and unfavourable in practice. If given unlimited resources, our current medicinal chemist toolkit should be able to satisfy a much larger area of chemical space. Nonetheless, cheap, efficient, and easily accessible routes are paramount to practically expand synthetically accessible chemical space. For the tool to be truly generalisable, more appropriate templates should be used. Alternatively, better, more appropriate, building blocks should be developed that allow for current templates to be used.

### S10 Classifying Useful Transformations

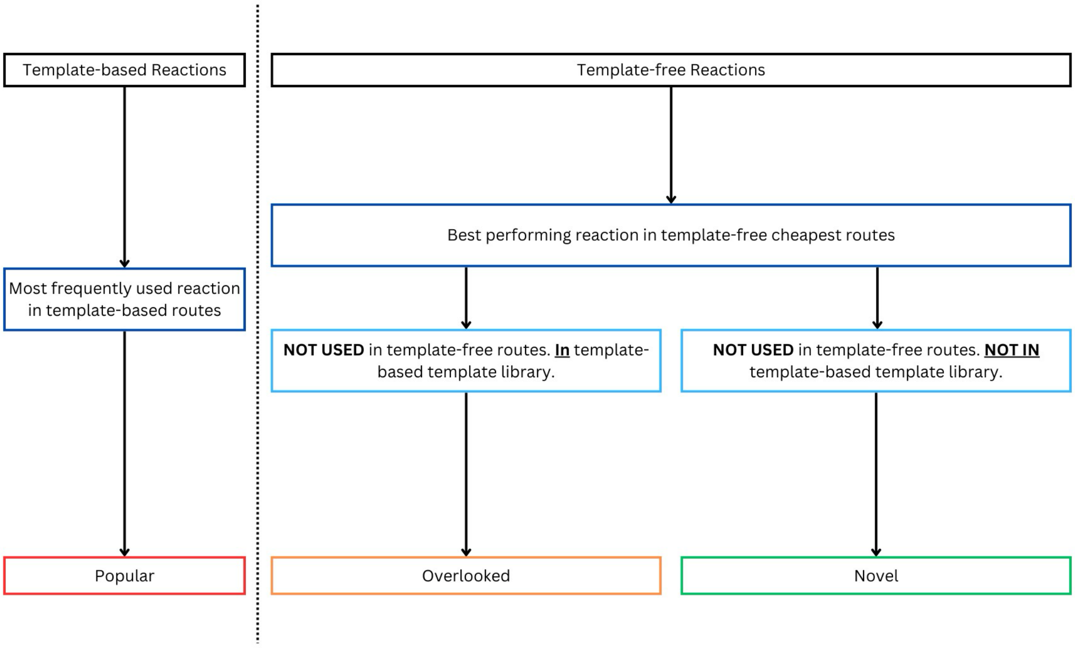


*Figure 6: Diagram detailing the classification of popular, overlooked and novel templates. Popular transformations are reactions frequently used in the most cost-effective TB routes. Overlooked reactions are those available to the TF method but not used. Novel reactions are neither used nor available to the TB method.*
